# P2G3 human monoclonal antibody neutralizes SARS-CoV-2 Omicron subvariants including BA.4 and BA.5 and Bebtelovimab escape mutants

**DOI:** 10.1101/2022.07.28.501852

**Authors:** Priscilla Turelli, Craig Fenwick, Charlène Raclot, Vanessa Genet, Giuseppe Pantaleo, Didier Trono

## Abstract

The rapid evolution of SARS-CoV-2 has led to a severe attrition of the pool of monoclonal antibodies still available for COVID-19 prophylaxis or treatment. Omicron subvariants notably escape most antibodies developed so far, with Bebtelovimab last amongst clinically approved therapeutic antibodies to display still good activity against all of them including the currently dominant BA.4/BA.5. We recently described P2G3, a broadly active SARS-CoV-2 monoclonal antibody, which targets a region of Spike partly overlapping with the site recognized by Bebtelovimab. Here, we reveal that P2G3 efficiently neutralizes SARS-CoV-2 omicron subvariants including BA.4/BA.5. We further demonstrate that P2G3 neutralizes Omicron BA.2 and BA.4 mutants escaping Bebtelovimab blockade, whereas the converse is not true.

**Funding:** EU COVICIS program; private foundation advised by CARIGEST SA.

---

Omicron was identified as a new SARS-CoV-2 (Severe Acute Respiratory Syndrome Coronavirus 2) variant in late 2021 and rapidly became dominant so as to account today for a large majority of new infections worldwide^1^. Omicron differs from previously prevalent SARS-CoV-2 variants at more than thirty amino acid positions within Spike, the protein responsible for mediating viral entry through recognition of the ACE2 receptor and targeted by SARS-CoV-2 neutralizing antibodies. Moreover, Omicron has been the background of rapid genetic drift, with the successive emergence of subvariants that rapidly unseat their predecessor, cycling through BA.1, BA.2 and as of July 2022, BA.4/5 (the Spike proteins of BA.4 and BA.5 are identical) along with BA.2 derivatives such as BA.2.12.1 and most recently BA.2.75, both classified as subvariants under monitoring by the WHO (https://www.who.int/en/activities/tracking-SARS-CoV-2-variants). Fueling this Omicron expansion are mutations that escape neutralization by most monoclonal antibodies (mAb) available for the treatment of COVID19 or by pre-Omicron convalescent sera. Accordingly, Omicron variants readily cause infections in individuals who have been either previously infected with other SARS-CoV-2 variants or subjected to full vaccination regimens, all current vaccines being derived from the original SARS-CoV-2 strain^2–7^.

Likely due to a combination of these immunological features and of variant-specific virological properties, Omicron is highly contagious yet generally causes milder symptoms compared with other SARS-CoV-2 variants including its immediate predecessor Delta^8,9^. This has led to the hope that the sweeping propagation of Omicron in the world population might contribute to attenuating the COVID19 pandemics by inducing, at a low cost for global health, robust levels of protective immunity to previously vaccinated and non-vaccinated populations alike. However, Omicron-induced immunity has turned out to be weak and short-lived, crushing this optimistic projection^10-11^. The continuous spread of immunity-evading Omicron subvariants thus represents an unabated threat, in particular for individuals at risk of developing serious SARS-CoV-2-induced disease due to immune dysfunctions or other underlying pathologies.

We recently described two human monoclonal antibodies with potent neutralizing activity against a wide array of SARS-CoV-2 variants^12, 13^. The first, P5C3, is a class 1 mAb recognizing the viral Spike protein in the up configuration through a region overlapping by ∼70% with the receptor binding domain (RBD). P5C3 displays high levels of neutralizing activity against most early SARS-CoV-2 variants including Alpha and Delta as well as against Omicron BA.1 and BA.2^12, 13^. The second of these mAb, P2G3, is a class 3 monoclonal that binds Spike in both the up and down configurations, and is endowed with particularly high neutralizing activity against Omicron BA.1 and BA.2^13^. Cryo-EM studies further indicate that up to three P2G3 and one P5C3 molecules can simultaneously bind the Omicron Spike trimer. Importantly, by growing virus in the presence of increasing concentrations of antibody, we could identify mutants capable of escaping either P5C3 or P2G3, yet we observed that i) the corresponding mutations were extremely rare in the wild, ii) the mutant viruses had a reduced infectivity and iii) P2G3 or P5C3 cross-neutralized each other’s escape mutants. Further encouraged by *in vivo* experiments demonstrating that P2G3, alone or in combination with P5C3, conferred complete prophylactic or therapeutic protection in a non-human primate model of Omicron infection^13^, we examined the activity of these two mAbs against the latest Omicron subvariants and compared their performance with that of Bebtelovimab, the last amongst clinically approved therapeutic antibodies still to display strong activity against Omicron BA.4/5^10,14,15^.

To measure variant-specific neutralizing antibodies, we first used a cell-free Spike-ACE2 binding interference assay, the results of which we previously demonstrated to correlate faithfully with those of cell-based live virus neutralization systems^16^. We used trimeric Spike proteins derived from past and currently circulating variants or subvariants including the most recent Omicron BA.1, BA.2, BA.2.12.1 and BA.4/5 strains (Table 1 and Figure 1). We found P2G3 to compete with ACE2 for the binding of all these Spike derivatives with equal efficiencies, exhibiting a profile similar to Bebtelovimab. P5C3 prevented ACE2 binding to Spike proteins from all tested variants except for BA.4/5, a results that was anticipated since in this pair of subvariants a valine replaces phenylalanine 486, a residue located at the interface between Omicron Spike and the P5C3 Fab^13^.

**Table 1:** Spike mutations present in indicated variants, compared with Wuhan strain

|  |  |
| --- | --- |
| **D614G** | D614G |
| **Alpha**<br>**B.1.1.7** | Δ69-70, Δ144, N501Y, A570D, D614G, P681H, T716I, S982A, D1118H |
| **Beta**<br>**B.1.351** | L18F, D80A, D215G, Δ242-244, R246I, K417N, E484K, N501Y, D614G, A701V |
| **Gamma**<br>**P.1** | L18F, T20N, P26S, D138Y, R190S, K417T, E484K, N501Y, D614G, H655Y, T1027I, V1176F |
| **Delta**<br>**B.1.617.2** | T19R, Δ156-157, R158G, L452R, T478K, D614G, P681R, D950N |
| **Delta**<br>**AY.4.2** | T19R, T95I, Y145H, Δ156-157, R158G, A222V, L452R, T478K, D614G, P681R, D950N |
| **Kappa**<br>**B.1.617.1** | E154K, L452R, E484Q, D614G, P681R, Q1071H |
| **Lambda**<br>**C.37** | G75V, T76I, R246N, del247-253, L452Q, F490S, D614G, T859N |
| **Iota**<br>**B.1.526** | L5F, T95I, D253G, E484K, D614G, A701V |
| **Eta**<br>**B.1.525** | Q52R, del69-70, E484K, Q677H, F888L |
| **Omicron**<br>**BA.1** | A67V, Δ69-70, T95I, G142D, Δ143-145, Δ211, L212I, ins214EPE, G339D, S371L, S373P, S375F, K417N, N440K, G446S, S477N, T478K, E484A, Q493R, G496S, Q498R, N501Y, Y505H, T547K, D614G, H655Y, N679K, P681H, N764K, D796Y, N856K, Q954H, N969K, L981F |
| **Omicron**<br>**BA.1.1** | A67V, Δ69-70, T95I, G142D, Δ143-145, Δ211, L212I, ins214EPE, G339D, R346K, S371L, S373P, S375F, K417N, N440K, G446S, S477N, T478K, E484A, Q493R, G496S, Q498R, N501Y, Y505H, T547K, D614G, H655Y, N679K, P681H, N764K, D796Y, N856K, Q954H, N969K, L981F |
| **Omicron**<br>**BA.2** | T19I, Δ24-26, A27S, G142D, V213G, G339D, S371F, S373P, S375F, T376A, D405N, R408S, K417N, N440K, S477N, T478K, E484A, Q493R, Q498R, N501Y, Y505H, D614G, H655Y, N679K, P681H, N764K, D796Y, Q954H, N969K |
| **Omicron**<br>**BA.2 12.1** | T19I, Δ24-26, A27S, G142D, V213G, G339D, S371F, S373P, S375F, T376A, D405N, R408S, K417N, N440K, L452Q, S477N, T478K, E484A, Q493R, Q498R, N501Y, Y505H, D614G, H655Y, N679K, P681H, S704L, N764K, D796Y, Q954H, N969K |
| **Omicron**<br>**BA.4 and**<br>**BA.5** | T19I, Δ24-26, A27S, Δ69-70, G142D, V213G, G339D, S371F, S373P, S375F, T376A, D405N, R408S, K417N, N440K, L452R, S477N, T478K, E484A, F486V, Q498R, N501Y, Y505H, D614G, H655Y, N679K, P681H, N764K, D796Y, Q954H, N969K |

**Figure 1.**
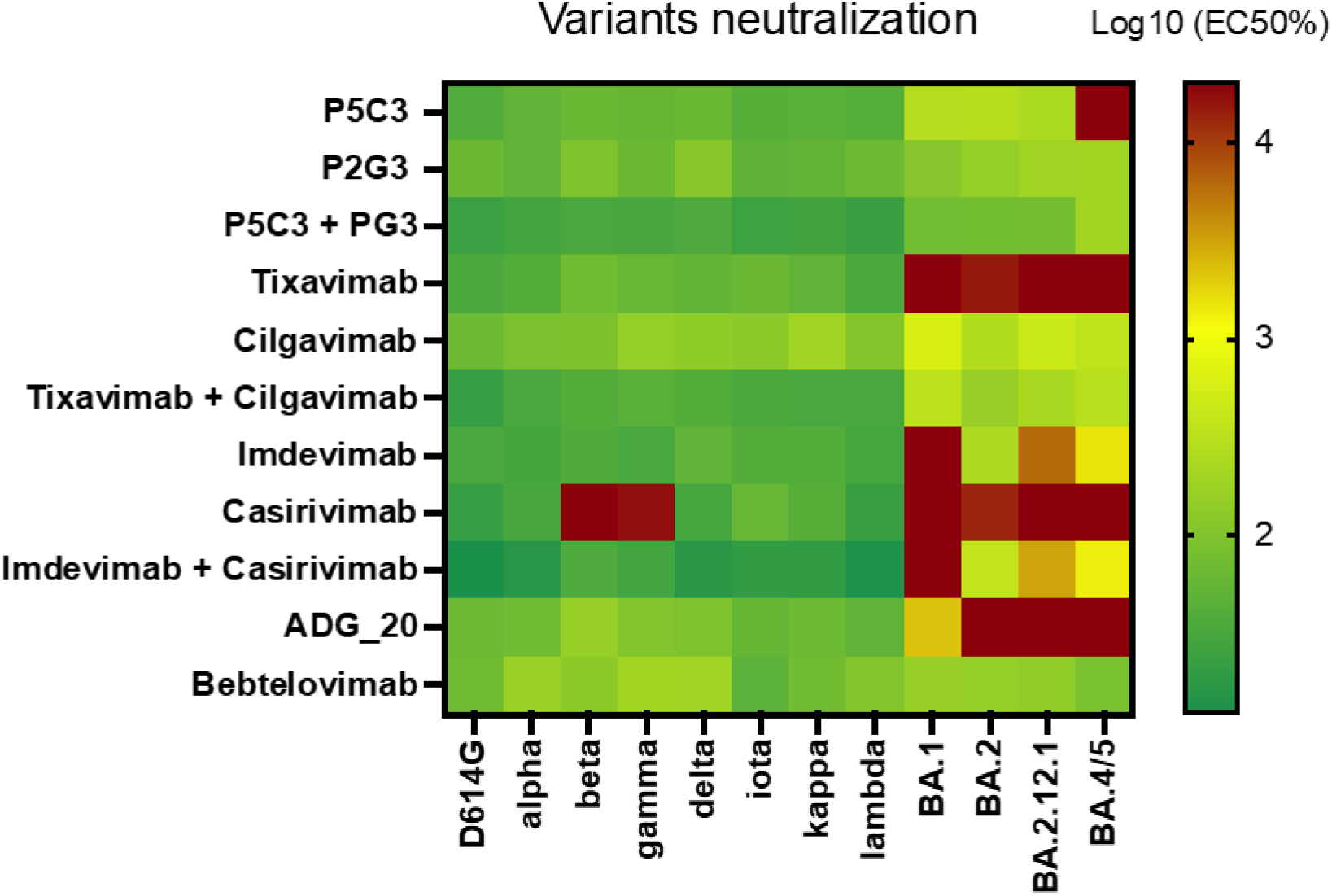
Heatmap representation of EC50 obtained in cell-free surrogate neutralization assay^16^ with indicated Spike proteins and antibodies.

Activities in the cell-free Spike-ACE2 interaction assay were consistent with results previously obtained for D614G, Alpha, Beta, Gamma, Delta, BA.1 and BA.2 strains in cell-based neutralization assays with the corresponding live viruses^13^. We thus also used replication-competent BA.4 and BA.5 infection of Vero cells with measurement of cytopathic effect by plaque counting (Figure 2 and Table 2). It confirmed that P2G3 and Bebtelovimab neutralized Omicron BA.4 and BA.5 with comparable efficiencies, with IC50 and IC80 <20 and <30 ng/ml, respectively, for both antibodies, that is, between 8 and 15 times lower than measured for AZD8895/1061 (Tixagevimab/Cigalvimab) or RGN10933/10987 (Casirivimab/Imdevimab) combinations, and 30 to >100 times lower than obtained for ADG-2 and Sotrovimab, respectively.

**Figure 2.**
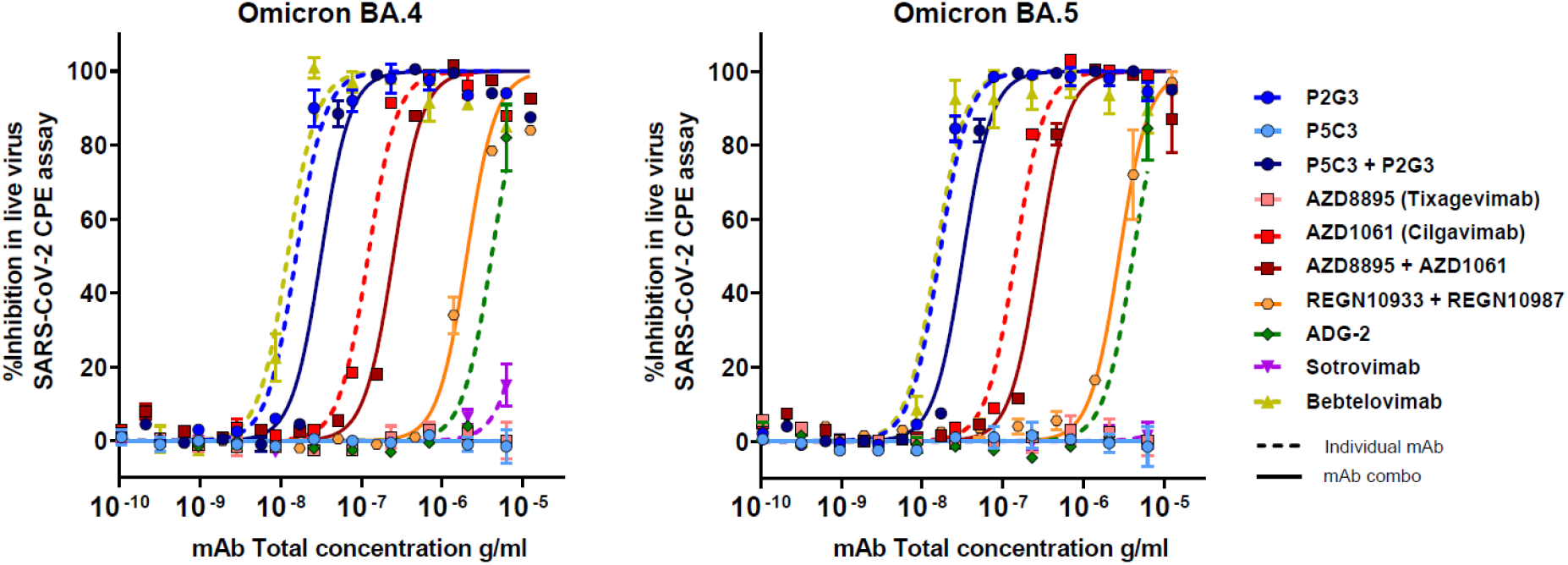
Neutralization activity of indicated antibodies measured in a live SARS-CoV-2 infectious virus cytopathic effect assay (CPE). Replication-competent BA.4 and BA.5 viruses were used to infect Vero E6 *in vitro* in the absence and presence of increasing concentrations of the indicated mAbs, individually or in combinations.

**Table 2:** IC50 and IC80 measured for indicated antibodies with BA.4 and BA.5 in live virus, cytopathic effect-based neutralization assay.

| Individual and combination mAbs | Omicron BA.4 |  | Omicron BA.5 |  |
| --- | --- | --- | --- | --- |
|  | IC50 | IC80 | IC50 | IC80 |
| P2G3 | 0.015 | 0.027 | 0.016 | 0.029 |
| P5C3 | >20 | >20 | >20 | >20 |
| P2G3 + P5C3 | 0.031 | 0.056 | 0.032 | 0.057 |
| AZD1061 | 0.121 | 0.216 | 0.143 | 0.255 |
| AZD8895 | >20 | >20 | >20 | >20 |
| AZD1061 + AZD8895 | 0.249 | 0.444 | 0.281 | 0.5 |
| REGN10933 + REGN10987 | 2 | 3.6 | 2.8 | 4.9 |
| ADG-2 | 4.1 | 7.3 | 4.1 | 7.3 |
| Sotrovimab | >20 | >20 | >20 | >20 |
| Bebtelovimab | 0.012 | 0.022 | 0.015 | 0.026 |

| IC<sub>50</sub> or IC<sub>80</sub> (μg/ml) |
| --- |
| <0.030 |
| 0.030 to 0.080 |
| 0.080 to 0.200 |
| 0.200 to 0.400 |
| 0.400 to 1.0 |
| >1.0 |

We then proceeded with viral resistance studies through the *in vitro* selection of P2G3 and Bebtelovimab BA.2 and BA.4 escape mutants as previously described^13^. Although no BA2 escapee was obtained in these experiments with either P2G3 or P5C3 alone or in combination, mutations at position 444 (K444E or K444M) appeared in sequences of BA.4 viruses resistant to these antibodies. Since P5C3 does not neutralize the BA.4 subvariant and since we previously documented the K444T P2G3 escape mutation in the Delta background^13^, it follows that the loss of K444 abrogates P2G3-mediated neutralization of BA.4. Resistance studies with Bebtelovimab selected the K444T and V445A escape mutations for BA.2, and the V445F substitution for BA.4 in our cell culture system. The distinct escapees profile of P2G3 and Bebtelovimab prompted us to ask whether they would cross-neutralize each other’s resistant mutants. For this, we used both live virus assays and lentivector particles pseudotyped with Delta and Omicron Spike derivatives.

With live virus, we found that Bebtelovimab could not neutralize P2G3-resistant BA.4, consistent with its dependance on the presence of K444 for efficient blockade. In contrast, P2G3 did neutralize the viral pool of Bebtelovimab escapees, suggesting that it was less sensitive to the V445F mutation (Figure 3).

**Figure 3.**
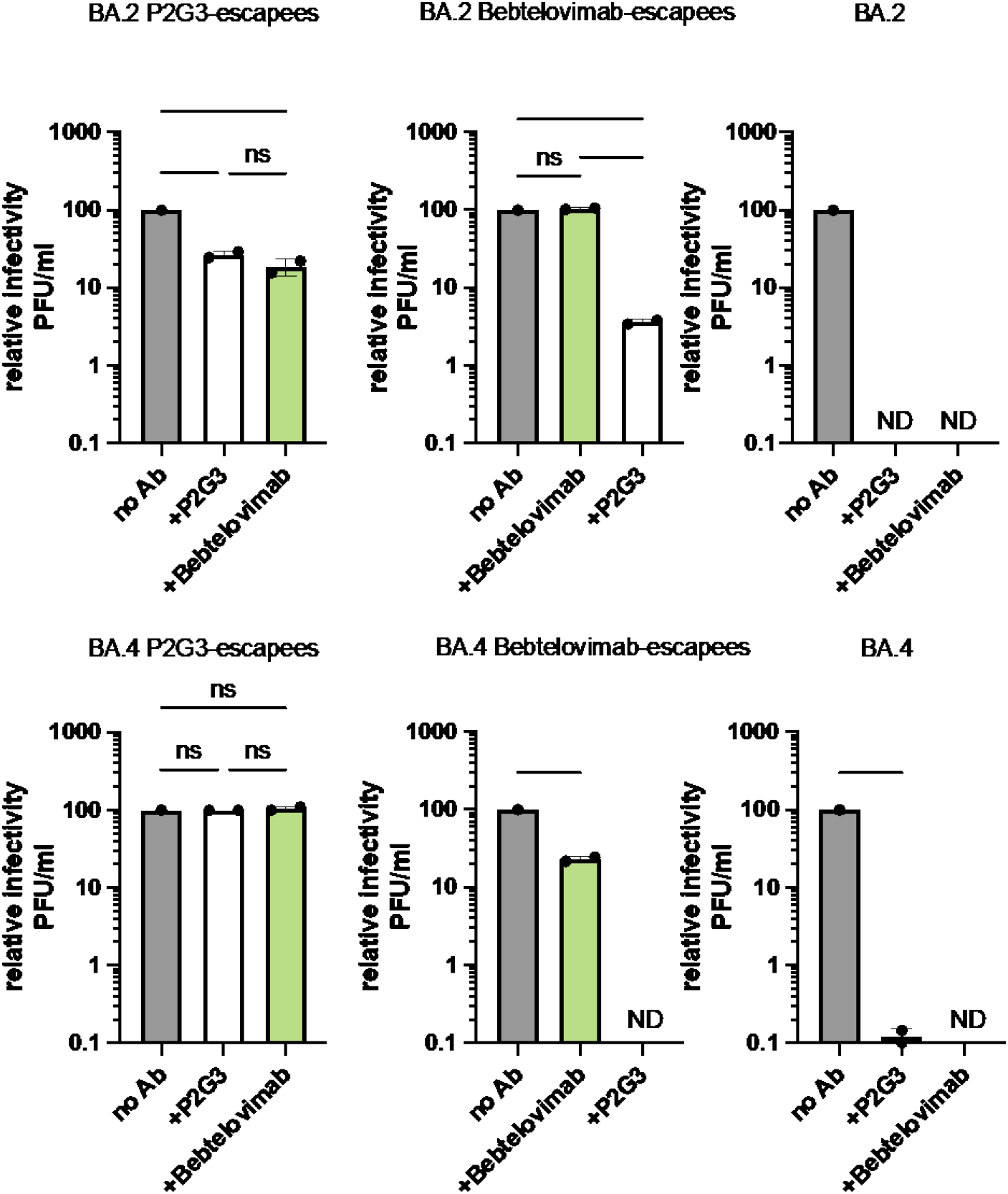
Infectious score of indicated viruses in the absence (no Ab) or presence of inhibitory concentrations of P2G3 (+P2G3) or Bebtelovimab (+Lycov 1404). Number of plaque-forming units (PFU) per ml obtained in the absence of antibody is given in each case a value of 100.

We explored further the activity of P2G3, P5C3 and Bebtelovimab on each other’s resistant mutants by using the cell-free Spike-ACE2 binding competition assay with a series of wild-type and mutated Spike derivatives with substitutions identified as mediating escape (Fig. 4). The results confirmed that the K444T and to a lesser extent the V445G mutations conferred P2G3 resistance in the Delta background, while these amino acid substitutions had less impact when introduced in the Omicron Spike protein. With Bebtelovimab, resistance was also confirmed for the K444T mutation, and was more pronounced than with P2G3 for the V445G substitution. Importantly, while the BA.1 strain harboring a G446S mutation stays sensitive to Bebtelovimab, we noticed that a BA.1 or BA.2 Spike harboring a G446V substitution was resistant to this mAb but more sensitive to P2G3. In contrast, a G446A substitution did not abrogate Bebtelovimab-mediated blockade, indicating that it is the introduction of a V at this position, nor the mere loss of a G, that precludes neutralization by this mAb.

**Figure 4.**
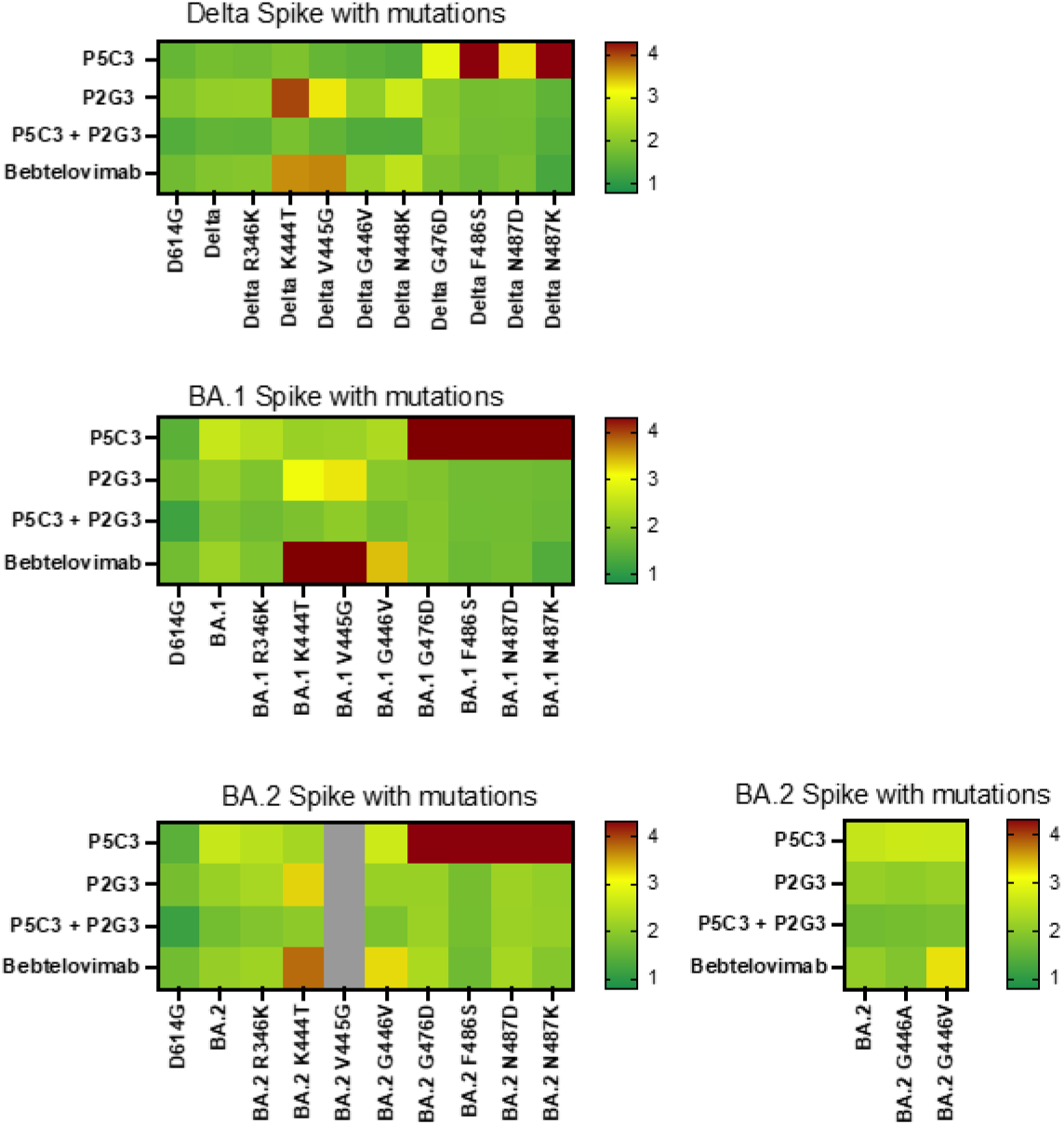
Heatmap representation of neutralization activity of P2G3, P5C3 and Bebtelovimab against indicated Spike derivatives in cell-free ACE2-Spike binding assay.

We went on to test cross-neutralization of P2G3/Bebtelovimab escape mutants using Spike-pseudotyped lentivector particles. In a Delta Spike background, mutations K444T, V445G and G446V conferred resistance to P2G3 and Bebtelovimab, but Spike derivatives harboring these changes where all blocked by P5C3 (Fig 5, a-e). In the Omicron BA.4 background, either K444T or V445G suppressed P2G3 and Bebtelovimab action, whereas G446A had no effect. Most interestingly, mutation F499H, reported by the manufacturer as allowing Bebtelovimab escape, indeed conferred resistance to this mAb, but had no impact on P2G3-mediated neutralization (Fig. 5, f-k).

**Figure 5.**
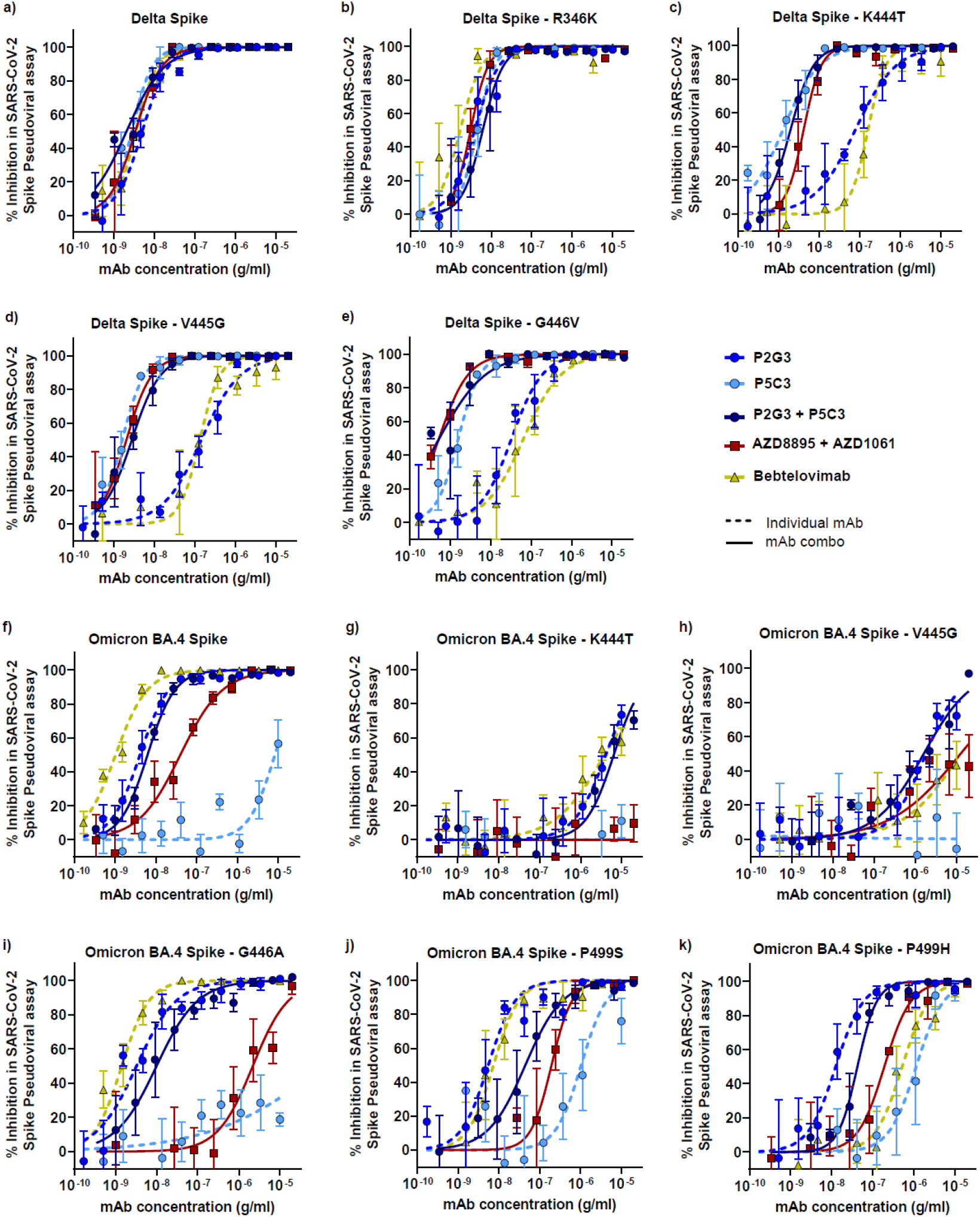
Neutralization of lentiviral particles pseudotyped with indicated Delta (a-e) or Omicron BA.4 (f-k) Spike derivatives in a 293T-ACE2 infection assay. Antibody cocktails representing a 1:1 mix of each mAb to give the indicated total mAb concentration.

We next compared the RBD binding epitopes for P2G3 and Bebtelovimab obtained from cryo-EM structures for the respective Fab’s bound to Spike trimer^13, 17^ (Fig 6). Residues K444 and V445 formed direct contacts with both mAbs and is consistent with the importance of these residues for maintained activity in both Spike-ACE2 and pseudoviral assays. The G446 residue is within the contact regions for both mAbs, but is more peripheral to the P2G3 binding epitope which may explain the enhanced sensitivity of Bebtelovimab to the G446V and G446F substitutions. Furthermore, the selective resistance to Bebtelovimab induced by the P499 mutations is consistent with our structural evaluation, which reveals this amino acid as a central point of contact for Bebtelovimab but not P2G3.

**Figure 6.**
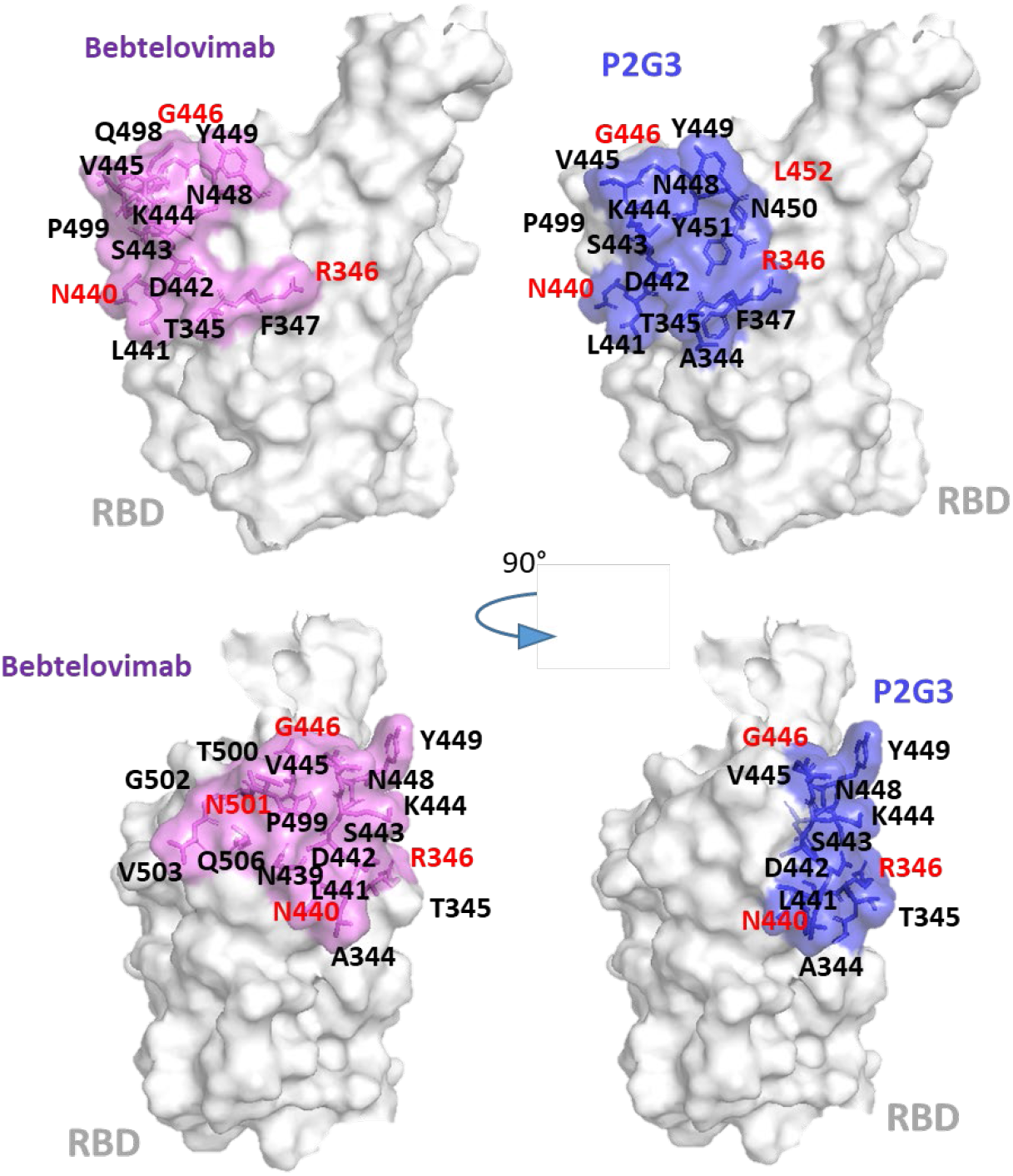
Comparison of the contact points of P2G3 and Bebtelovimab with the Spike RBD protein. Bottom set of RBD models are rotated 90° relative to the top panels to display the full extent of the antibody binding epitopes. Amino acid residues in red represent mutations present in Omicron spike variants that are proximal to the binding epitopes for the two antibodies. N440K, N501Y mutations are present in all Omicron variants, G446S in BA.1, R346K in BA.1.1, L452Q in BA.2.12.1 and L452R in BA.4/5.

The *in vitro* selection of viral escapees is an important tool to assess potential resistance mutations to neutralizing antibody. However, viral fitness and infectivity in a natural setting will ultimately define the likelihood these mutation will emerge. To help define these *in vivo* fitness properties, escapees to P2G3 and Bebtelovimab were matched to their prevalence in the GISAID sequence database (https://www.gisaid.org/hcov19-variants/). Substitutions at K444 conferring resistance to both mAbs were extremely rare with K444T, K444E and K444M identified in 0.0039%, 0.0005% and 0.0022% respectively of the 12’086’753 available sequences as of July 2022. Prevalence of the V445G was also rare at 0.00029% while G446V mutation was detected at comparatively high levels in being detected in 0.1205% of sequences. The P499H escapee with selective resistance to Bebtelovimab is rare at 0.0011% sequence prevalence, however the P499R substitution that is reported to confer an equal degree of resistance (https://www.fda.gov/media/156152/download) was detected in 0.015% of sequenced worldwide viral isolates.

Altogether, these results reveal that P2G3 is a highly efficient monoclonal antibody that is capable of neutralizing all currently prevalent Omicron subvariants, notably BA.4/BA.5, and appears to display an advantageously narrower range of escapees compared with Bebtelovimab. Considering the demonstrated in vivo efficacy of P2G3, alone or in combination with P5C3, for both the prevention and the treatment of Omicron infection in the non-human primate model and the availability of both of these antibodies as extended half-life derivatives, these results strongly support the use of P2G3/P5C3 for both prophylactic and therapeutic purposes, and warrant their upcoming testing in a phase I clinical trial.

## METHODS

### Spike proteins production and purification, S^3^-cell free neutralization assay

Trimeric-Spike variants production and purification and Spike protein beads coupling were performed as previously described^15^. Neutralization assays were done in 96-well plates as previously described^15^, with 10 multiplexed Spike variants.

### SARS-CoV-2 live virus stocks

All biosafety level 3 procedures were approved by the Swiss Federal Office of Public Health (authorization A202952/3). The SARS-CoV2 D614G (EPI_ISL_414019), Alpha (EPI_ISL_2131446), Beta (EPI_ISL_981782), Gamma (EPI_ISL_981707), Delta (EPI_ISL_1811202) and Omicron BA.1 (EPI_ISL_7605546), BA.2 (EPI_ISL_8680372), BA.4 (EPI_ISL_12268495.2) and BA.5 (EPI_ISL_12268493.2) early passage isolates were obtained and sequenced at the Geneva University Hospitals. Viral stocks were prepared with the early isolates in EPISERF on VeroE6 or Calu-3 cells, aliquoted, frozen and titrated on VeroE6 cells.

### SARS-CoV-2 live virus cell based cytopathic effect neutralization assay

Neutralization assay was performed as previously described^13^ except that serial sera dilutions were done in EPISERF instead of DMEM 2% FCS. In brief, 3,000 plaque forming units per well of viral isolate were pre-incubated for 1 hour at 37°C with 1:3 serial dilutions of sera prepared in duplicates in 96-well plates. The mix was then applied on VeroE6 cells and cytopathic effect detected 48hours later with Crystal violet staining.

### Selection of escape mutants

Omicron BA.2 and BA.4 viral isolates were used to infect 293T cells over expressing ACE2 and TMPRSS2 (MOI of 0.2) each in duplicates in presence of suboptimal concentrations of antibodies (0.25ng/ml) as previously described^13^. Briefly, supernatants were collected, diluted 40-fold and used to infect fresh cells for two more passages for four days and in the same conditions (P1 to P3). Putative viral escapees were further selected by serial passages of 2-fold diluted supernatants pre-incubated 1hrs at 37°C with high concentrations of antibodies (2.5 μg/ml or 0.625 μg/ml, each tested in duplicates). Viral RNA extracted from supernatants collected at each passage was deep-sequenced and P5 viral supernatant used for neutralization assays. Virus produced in absence of mAb was collected and treated the same way in parallel to control for appearance of mutations due to cell culture conditions.

### Cross neutralization of viral escapees

Supernatants collected from selection experiments performed in duplicates with 2,5µg/ml of antibodies were pre-treated with 10µg/ml of antibodies for 1hrs at 37°C and applied on VeroE6 cells in presence of antibodies and Avicell 0.4% overlay. Cells were stained 3 days later with crystal violet and plaques were counted.

## ACKNOWLEDGMENTS

We thank Florence Pojer and the Protein production and structure Core facility at EPFL for the Spike variants production, Meriem Bekliz and the Virology laboratory of Geneva University Hospital for the Omicron RNA samples and variant isolates collection.

